# Predicting long-term collective animal behavior with deep learning

**DOI:** 10.1101/2023.02.15.528318

**Authors:** Vaios Papaspyros, Ramón Escobedo, Alexandre Alahi, Guy Theraulaz, Clément Sire, Francesco Mondada

## Abstract

Deciphering the social interactions that govern collective behavior in animal societies has greatly benefited from advancements in modern computing. Computational models diverge into two kinds of approaches: analytical models and machine learning models. This work introduces a deep learning model for social interactions in the fish species *Hemigrammus rhodostomus*, and compares its results to experiments and to the results of a state-of-the-art analytical model. To that end, we propose a systematic methodology to assess the faithfulness of a model, based on the introduction of a set of stringent observables. We demonstrate that machine learning models of social interactions can directly compete against their analytical counterparts. Moreover, this work demonstrates the need for consistent validation across different timescales and highlights which design aspects critically enables our deep learning approach to capture both short- and long-term dynamics. We also show that this approach is scalable to other fish species.

## 1 Introduction

Collective behavior in animal groups is a very active field of research, studying the fundamental mechanisms by which individuals coordinate their actions [1–3] and self-organize [4, 5]. One of the most common forms of collective behavior can be observed in schools of fish and flocks of birds that have the ability to coordinate their movements to collectively escape predator attacks or improve their foraging efficiency [6, 7]. This coordination at the group level mainly results from the social interactions between individuals. Important steps to understand these collective phenomena consist in characterizing these interactions and understanding the way individuals integrate interactions with other group members [8–11].

New tracking techniques and tools for behavioral analysis have been developed that have greatly improved the quality of collective motion data [12–18]. In particular, advances in computing have allowed the development of computationally demanding data-oriented model generation techniques [11, 19–23] and the subsequent simulation of biological models [24]. This has resulted in more realistic models that attempt to recover the social interactions that govern collective behaviors. Yet, the bottleneck with most of these approaches is that they rely on demanding and laborious mathematical work to obtain the interactions from experimental data.

An alternative to such analytical models is to exploit machine learning (ML) techniques and let an algorithm learn the interactions directly from data. The know-how required to use these techniques is different from the one needed to design analytical models. Nevertheless, the structure of ML algorithms, here a neural network, has an impact on the modeling performance, and requires specific expertise [25]. Once the architecture of an ML algorithm is set, ML can often process data for different species without structural adaptation, and generate new models quickly. This is very different from analytical models, where each new species requires redefining the model nearly from scratch. The downside of this flexibility is that ML models are usually less explainable (“black box”). Yet, recent ML algorithms provide higher-level information mappable to more tangible formats, such as force maps, which show the strength and direction of behavioral changes experienced by an individual when interacting with other individuals in a moving group [22, 23]. Despite their limited explainability, ML algorithms require only few biological assumptions. They offer an almost hypothesis-free procedure [26] that can even outperform human experts in detecting subtle patterns [27], making ML a very appealing alternative or complement to analytical models.

For both analytical and ML models, several studies evaluate models over *short timescales* and through instantaneous quantities such as speed, acceleration, distance and angle to objects [21, 28], or by measuring the error between predictions and ground truth [22, 29, 30]. Only more recently, long timescales have also been considered [20]. However, a model that performs well at short timescales compared to experiment does not necessarily perform well at long timescales. This is especially true for models that try to reproduce complex collective phenomena in living systems. To our knowledge, the predictive capacity of ML models in this context has not been evaluated over both short *and* long timescales, that is, their ability to generate synthetic data that replicate the outcomes of social interactions over both timescales.

Here, we demonstrate that ML models can generate realistic synthetic data with minimal biological assumptions, and that they allow to accelerate and generalize the process of collective behavior modeling. More specifically, we present a social interaction model using a deep neural network that captures both the short- and long-term dynamics observed in schooling fish. We apply our approach to pairs of rummy-nose tetra (*Hemigrammus rhodostomus*) swimming in a circular tank, and show that it can be also applied to fish species with similar burst-and-coast swimming (zebrafish; *Danio rerio*). Our ML model is benchmarked against the state-of-the-art analytical model for this species [31], showing that it performs as well as the latter, even for very subtle quantities measured in the experiments. Moreover, we also introduce a systematic methodology to stringently test the results of an analytical or ML model against experiment, at different timescales, and in the context of animal collective motion.

## 2 Results

When fish swim in a circular tank (here, of radius *R* = 25 cm), they interact with each other and with the tank wall. The resulting collective dynamics can be finely characterized by exploiting the 9 observables introduced and described in the Methods Section. As explained there, these observables probe 1) the instantaneous individual behavior, 2) the instantaneous collective behavior, and 3) the temporal correlations of the dynamics.

Hereafter, three 16-hour trajectory datasets are analyzed: the first one corresponds to pairs of *H. rhodostomus* in our experiment, the second one to the Analytical Burst-and-Coast model (ABC), and the third one to the Deep Learning Interaction model (DLI). Supplementary Video 1 shows typical trajectories for these three conditions. The aim of this section is to quantitatively validate the qualitative agreement observed in this video.

### 2.1 Quantification of the instantaneous individual behavior

The individual fish behavior is characterized by three observables: the probability distribution function (PDF) of the speed *V*, of the distance to the wall *r*_w_, and of the angle of incidence to the wall *θ*_w_. When swimming in pairs, fish tend to adopt a typical speed of about 7cm/s (see the peak of the PDF in Fig. 1A), but can also produce high speeds up to 25–30m/s. Both fish remain close to the wall of the tank (a consequence of the fish burst-and-coast swimming mode [11]), the leader being closer to the wall (typically, at about 0.5 BL) than the follower (at about 1.2 BL; see Fig. 1B). This feature is due to the follower fish trying to catch up the leader fish by taking a shortcut while taking the turn. Moreover, fish spend most of the time almost parallel to the wall: see the peaks of both PDFs at *θ*_w_ ≈ ±90° in Fig. 1C. A slight asymmetry is observed in the PDF of *θ*_w_, showing that, in the experiments, fish have turned more frequently in the counter-clockwise direction. Values of the mean and the standard deviation of the PDFs presented in this section are given in Supplementary Tables 2–4.

**Fig. 1:**
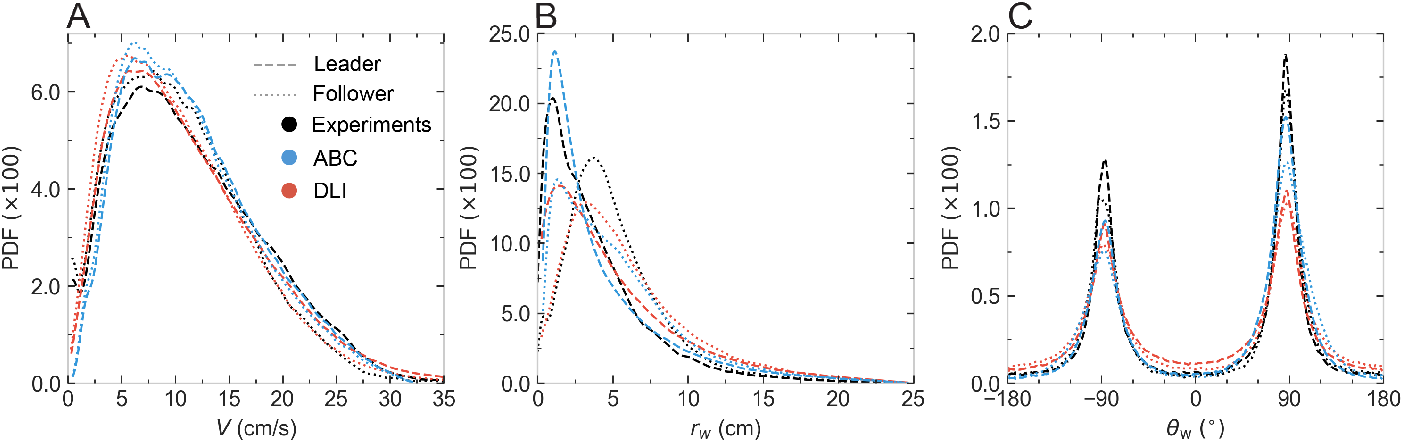
Probability density functions (PDF) of observables characterizing individual behavior: **A** Speed *V*, **B** distance to the wall *r*_w_, and **C** angle of incidence to the wall *θ*_w_. Black lines: experimental fish data. Blue lines: agents of the Analytical Burst-and-Coast model (ABC). Red lines: agents of the Deep Learning Interaction model (DLI). Dashed lines: geometrical leader; dotted lines: geometrical follower.

Both ABC and DLI models produce agents that move at the same mean speed as fish in the experiments, and Fig. 1A shows that the speed PDF for both models are in excellent agreement with the one observed in real fish. Moreover, the agents of the ABC model are as close to the wall and as parallel to it as fish are. The PDF of the ABC leader is in good agreement with that of the fish leader (Fig. 1B). However, the PDF for the ABC follower has a peak at about the same distance to the wall as that of the leader, while the corresponding peaks are more separated for real fish. Yet, the PDF for the ABC follower is broader than for the leader, showing that the ABC follower tends to be farther from the wall than the leader, as observed for real fish. For the DLI model, the peaks of both leader and follower PDFs are at about the same position as for real fish, although their height is smaller than for fish, meaning that DLI-agents tend to explore more frequently the interior of the tank (observe the thicker tails of the PDF of *r*_w_ for the DLI model in Fig. 1B). Alignment with the wall is also well reproduced by both models (Fig. 1C), including the asymmetry in the direction of rotation around the tank: their peak at *θ*_w_ > 0 is higher than the one at *θ*_w_ < 0. As already seen in the PDF of *r*_w_, DLI-agents visit more often the interior of the tank, and are hence less aligned with the wall than the real fish and ABC agents. Note that the tendency of DLI-agents to rotate more frequently in the counterclockwise direction is learned from the training set, while this asymmetry has to be explicitly implemented in the ABC model, by introducing an asymmetric term in the analytical expression of the wall repulsion function. A closer look at Fig. 1C shows that fish actually follow the wall with a most likely angle of incidence |*θ*_w_| that is slightly smaller than 90°, a feature resulting from the burst-and-coast swimming mode inside a tank with positive curvature: fish are found more often going toward the wall than escaping it.

### 2.2 Quantification of the instantaneous collective behavior

*H. rhodostomus* is a social species, and Fig. 2A shows that the two fish remain most of the time close to each other, with the PDF of their distance *d_ij_* presenting a peak around *d_ij_* ≈ 7 cm ≈ 2 BL (mean and standard deviation of the PDFs presented in this section are given in Supplementary Tables 2–4). The fish have a strong tendency to align with each other, as shown in Fig. 2B, with the PDF of their relative heading *ϕ_ij_* being sharply peaked at 0°. In addition, the PDF of the viewing angle *ψ_ij_* show that the fish are swimming one behind the other rather than side-by-side. This is illustrated in Fig. 2C by the sharp difference in the PDF of the viewing angle for the leader and the follower. The PDF of *ψ*_leader_ is peaked around ±160°, meaning that the follower fish is almost right behind the leader fish, but slightly shifted to the right or left. A slight left-right asymmetry in the PDF of the viewing angles is also visible, the follower being more frequently on the left side of the leader, a consequence of the fact that the fish in the experiment follow the wall by turning more often counterclockwise (Fig. 1C), with the follower swimming farther from the wall than the leader (Fig. 1B).

**Fig. 2:**
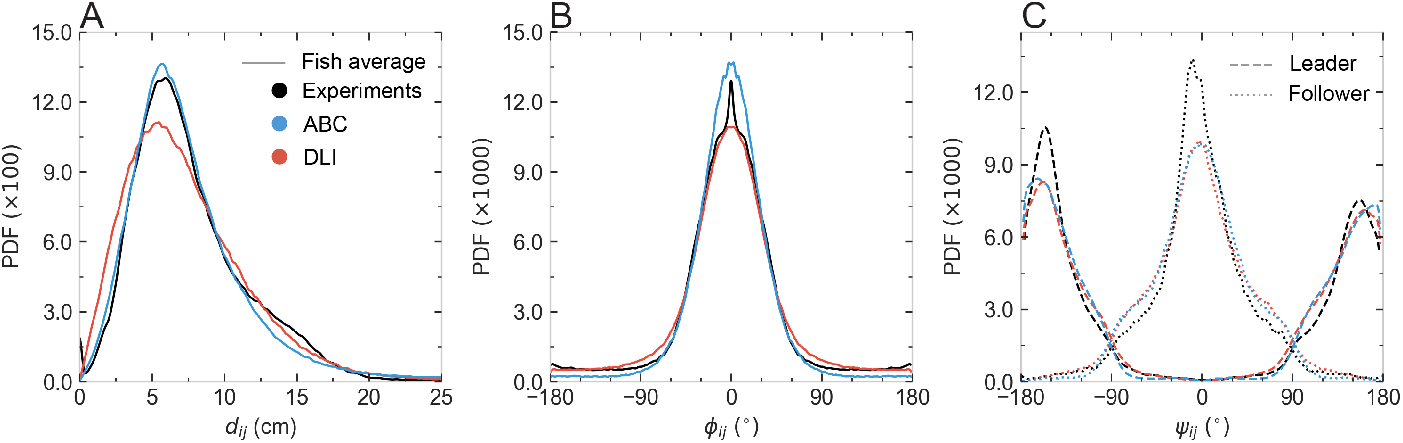
Probability density functions (PDF) of observables characterizing collective behavior: **A** Distance between individuals *d_ij_*, **B** difference in heading angles *ϕ_ij_*, and **C** angle of perception of the geometrical leader and follower *ψ_ij_*. Black lines: experimental fish data. Blue lines: agents of the Analytical Burst-and-Coast model (ABC). Red lines: agents of the Deep Learning Interaction model (DLI). Dashed lines : geometrical leader; dotted lines: geometrical follower (in **C**).

All these features are well reproduced by both models, with only some small quantitative deviations. The ABC model reproduces almost perfectly the experimental PDF of the distance between fish, whereas the PDF for the DLI model is only slightly wider and presents slightly more weight at very small distance than found for real fish or in the ABC model (Fig. 2A). The DLI model is in turn better than the ABC model at reproducing the PDF quantifying the alignment of the fish, the latter producing more weight near 0° than for real fish (Fig. 2B). Both models fail at reproducing the very small increase in the PDF at *ϕ_ij_* ≈ ±180°, which corresponds to sudden U-turns that real fish sometime perform. The PDF of the viewing angles for the leader and the follower (Fig. 2C) are also fairly reproduced by both models, including the slight left-right asymmetry observed in real fish, although the peak in the PDF at *ψ*_follower_ = 0° (and to a lesser extent at *ψ*_leader_ ≈ – 160°) is not quite as sharp as in the experiment.

### 2.3 Quantification of temporal correlations

Fig. 3 shows the 3 observables defined in equations (1–3) and probing the emerging temporal correlations in the system: the mean squared displacement *C_X_*(*t*), the velocity autocorrelation *C_V_*(*t*), and the autocorrelation of the angle of incidence to the wall *C*_*θ*_w__(*t*), as function of the time difference *t* between observations. The figure reveals that both models fail to fully reproduce quantitatively these very non-trivial observables, which indeed constitute the most challenging benchmark characterizing the correlations emerging from the fish behavior.

**Fig. 3:**
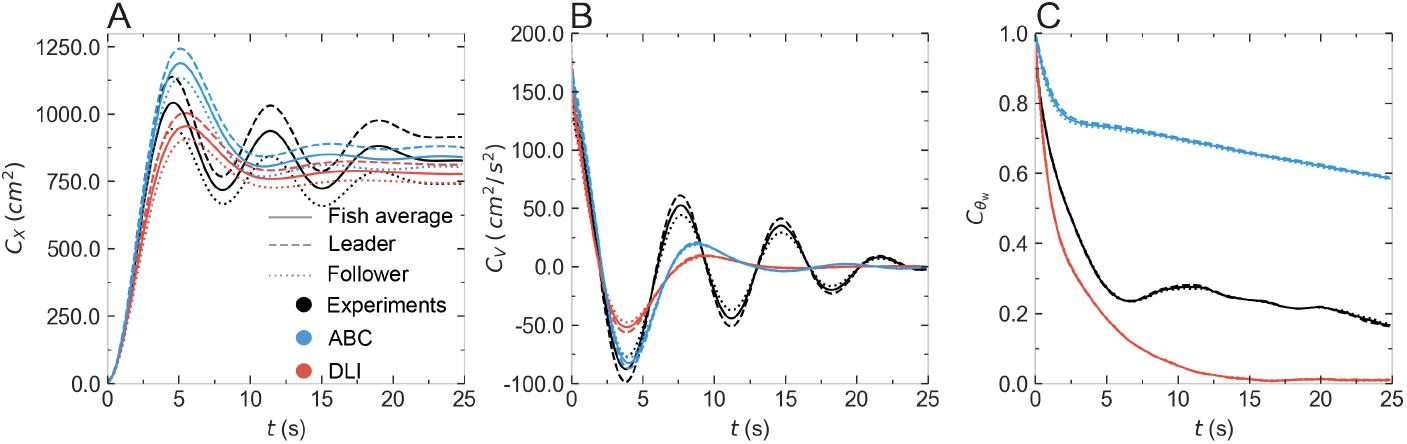
Observables quantifying temporal correlations in the system. **A** Mean squared displacement *C_X_*(*t*), **B** Velocity temporal autocorrelation *C_V_*(*t*), **C** Temporal correlations of the angle of incidence to the wall *C*_*θ*_w__ (*t*). Black lines: experimental fish data. Blue lines: agents of the Analytical Burst-and-Coast model (ABC). Red lines: agents of the Deep Learning Interaction model (DLI). Dashed lines: geometrical leader; dotted lines: geometrical follower; full lines: average over the 2 fish or agents.

Fish data present 3 distinct regimes: a quasi-ballistic regime at short timescale (*t* ≲ 1.5 s) where *C_X_*(*t*) ≈ 〈*v*^2^〉*t*^2^, followed by a second short diffusive regime (1.5 s ≲ *t* ≲ 5 s) where *C_X_*(*t*) ≈ *Dt*, which is limited by the finite size of the tank, ultimately leading to a third regime of saturation (*t* > 5 s) characterized by slowly damped oscillations due to the fact that fish are guided by the wall (Fig. 3A). Accordingly, the velocity correlation function starts from *C_V_*(*t* = 0) = 〈*v*^2^〉 at short time and also presents damped oscillations (Fig. 3B). The negative minima of the oscillations in *C_V_*(*t*) correspond to times when the focal fish is essentially at a position diametrically opposite to its position at the reference time *t* = 0, its velocity then being almost opposite to that at *t* = 0. Positive maxima similarly correspond to times when the fish returns to almost the same position it had at *t* = 0, with a similar velocity guided by the tank wall. Of course, these oscillations are damped as correlations are progressively lost, and the velocity correlation function *C_V_*(*t*) ultimately vanishes at large time *t* ≫ 20 s, due to the actual stochastic nature of the trajectories at this timescale (possible U-turns, or the fish randomly crossing the tank). Note that *C_X_* (*t*) is markedly different for the leader and follower fish, with a higher saturation value for the leader, which swims closer to the wall, as mentioned above.

The ABC model is able to fairly reproduce the short and intermediate regimes for *C_X_* (*t*) (Fig. 3A), as well as the position of its first peak, reached only slightly later than for fish (1s after). The ABC model also reproduces the experimental saturation value of *C_X_* (*t*) averaged over the two fish. As for the DLI model, its predictions are only slightly worse than that of the ABC model, due to the fact that the DLI agents are moving a bit farther to the wall compared to ABC agents and real fish. Yet, both models equally fail at producing more than one oscillation, and the correlations are damped faster compared to the experiment.

As for the velocity autocorrelation *C_V_*(*t*) (Fig. 3B), the ABC model reproduces almost perfectly the short and intermediate regimes and the position of the first negative minimum (hence, up to *t* = 6 s), while the DLI model underestimates the depth of this first minimum. But again, both models fail at reproducing the persistence of the correlations, producing a too fast damping of the oscillations (an effect slightly stronger in the DLI model).

Both models struggle at reproducing the correlation function *C*_*θ*_w__ (*t*) of the angle of incidence to the wall (Fig. 3C), where the fish curve first sharply decreases up to *t* = 6 s and then remains close to *C*_*θ*_w__ ≈ 0.2. The ABC model is clearly unable to reproduce both the decreasing range (clearly diverging before *t* = 2 s) and the correct saturation value (never falling below *C*_*θ*_w__ ≈ 0.6). As for the DLI model, it produces a slightly sharper decay of *C*_*θ*_w__ (*t*) than for real fish, up to *t* ≈ 6 s, but fails to reproduce the non-negligible remaining persistence of the correlation observed in fish for *t* > 7 s, with *C*_*θ*_w__ (*t*) in the DLI model decaying rapidly to zero. In fact, both models fail to reproduce the experimental *C*_*θ*_w__(*t*) for opposite reasons. The ABC model exhibits a too high persistence of the correlations of *θ*_w_ compared to real fish, presumably because real fish indeed often follow the wall but can also produce sharp U-turns, as observed in Fig. 1C. On the other hand, the failure of the DLI model in reproducing *C*_*θ*_w__ (*t*) stems from the fact that DLI agents move farther from the wall and cross through the tank more often than real fish and ABC agents (see the discussion of Fig. 1B above), hence leading to a too fast, and ultimately total, loss of correlation for *θ*_w_.

### 2.4 Complementary analyses

We also conducted three complementary tests of our approach. First, the DLI yielded better results in generating social interactions than a similarly purposed ANN (see Supplementary Figures 1 and 2, and Supplementary Video 2) for human trajectory forecasting [29, 30]. While this is expected, these results confirm that there exist models that do indeed capture the short-term dynamics without being able to reproduce the long-term dynamics. Secondly, we trained the DLI with data for pairs of zebrafish (*D. rerio*), and found that it yields good results for this species too, without any structural modification in its architecture (see Supplementary Figure 3). While acquiring a functional model of a new species’ interactions proved straightforward with the DLI, the same would not be generally true for analytical models.

We plan to exploit the DLI model to study groups with more than two fish, *without any retraining*. Indeed, *H. rhodostomus* [32], like many other group-living species [7], effectively only interact with a few influential neighbors, at a given time. Thus, for a given agent in a group of *N* > 2 agents, the DLI for *H. rhodostomus* should only retain the influence of typically the two agents leading to the highest acceleration [32, 33], as predicted by the DLI model. Supplementary Video 4 illustrates this procedure for *N* = 5 agents, resulting in a cohesive and aligned group, in qualitative agreement with experimental observation [32].

## 3 Discussion

Studying social interactions in animal groups is crucial to understand how complex collective behaviors emerge from individuals’ decision-making processes. Very recently, such interactions have been extensively investigated in the context of collective motion by exploiting classical computational modeling [11, 19, 20] and automated machine learning-based methods [22, 23]. Although ML algorithms have been shown to provide insight into the interactions of hundreds of individuals at short timescales [22, 23], their ability to reproduce the complex dynamics in animal groups at long timescales has not yet been assessed.

Here, we have presented a deep learning interaction model (DLI) which reproduces the behavior of fish swimming in pairs. We have also introduced the appropriate tools for its validation when compared to experimental results and when confronted with the state-of-the-art analytical model (ABC). In fact, our study establishes a systematic methodology to assess the long-term predictive power of a model (analytical or ML), by introducing a set of fine observables probing the individual and collective behavior of model agents, as well as the subtle correlations emerging in the system. These observables, which can be straightforwardly extended to groups of *N* > 2 agents, provide an extremely stringent test for any model aimed at producing realistic long-term trajectories mimicking that of actual animal groups. In particular, we consider that the usual validation of an ML model at a short timescale should be complemented by the type of long timescale analysis that we propose here, in order to fully assess its performance.

The DLI model closely reproduces the dynamics of real fish at both the individual (speed, distance to the wall, angle of incidence to the wall) and collective (distance between individuals, relative heading angle, angle of perception) levels during long simulations corresponding to more than 16 hours of fish swimming in a tank, hence successfully generating life-like interactions between agents. When compared to experiment, the ABC model and the DLI model essentially performs equally well. Notably, the DLI model better captures the most likely distance of the leader and follower from the wall. However, the DLI model is less accurate in reproducing the temporal correlations quantified by the mean-squared displacement and the velocity autocorrelation. Yet, both ABC and DLI models fail at capturing the temporal correlations of the angle of incidence to the wall, but for very different reasons.

Our study demonstrates two advantages of ML techniques: 1) they can drastically accelerate the generation of new models (as illustrated here for zebrafish), and 2) with minimal expertise in biology or modeling. This is especially useful in robotics, where models often act as behavioral controllers (*i.e*., trajectory generators) that guide the robot(s). Although there already exist many bio-hybrid experiments in the literature, most of them rely on simplified models for behavioral modulation [34–36], few of them exploit realistic models (analytical or ML) [28, 37], and, to our knowledge, none of them are tested in the long term in simulations or real-life. In this context, ML has the potential to benefit multidisciplinary studies, provided such techniques are thoroughly validated in simulations.

However, accelerating the production of collective behavior models with ML comes at a cost. Indeed, the DLI is a black-box model, and although it captures the subtle impact of social interactions between individuals, it is impossible to retrieve the interaction functions themselves. Some approaches partially address this issue by providing insight into how the network operates for specific sets of inputs [22, 23]. Yet, they still do not offer explicit interaction functions. On the other hand, analytical models supplemented by a procedure to reconstruct social interactions [11, 19] provide a concise and explicit description of the system in question. Moreover, varying the parameters of such models allows for investigating their relative impact on the dynamics, and to make predictions for various sets of these parameters [33]. This is not feasible with ML models, unless they are retrained or specifically structured to allow it.

In summary, this work shows that DLI-like models may now be considered as firm candidates to shed light on groundbreaking problems such as how social interactions take place and affect collective behavior in living groups. Yet, we have emphasized that social interaction models should be precisely tested at both short *and* long timescales. Future work includes the design of ANNs that provide additional information about the learned dynamics, possibly by exploiting symbolic regression algorithms [38, 39]. We also plan to study the extension of the DLI model to larger groups, in particular, in connection with our robotic platform [34–36]. We believe that this could improve our understanding of the mechanisms arising in collective behavior and allow for more precisely exploring and modulating them.

## 4 Methods

### 4.1 Experimental data

The trajectory data used in this study were originally published in [11] for *Hemigrammus rhodostomus* swimming either alone or in pairs in a 50 cm diameter circular tank [11]. This species is characterized by a burst-and-coast swimming mode, where the fish perform a succession of sudden short acceleration periods (kick), each followed by a longer gliding period almost in a straight line. The instant of the kicks, when heading changes take place, are assimilated to decision instants.

The dataset corresponds to 40 hours of video recordings at 25 Hz. Fish are tracked with idTracker [16], an image analysis software which extracts the 2D trajectories of all individuals. Occasionally, the tracking algorithm is temporarily unable to report positions accurately. This can be due to small fluctuations in lighting conditions, fish standing still or moving at very low speed, fish swimming very close to the surface, to the border, or to each other. These instances are corrected using several filtering processes. Since our analyses focus on social interactions, we remove the periods during which fish are inactive. Fish body length (BL) is of about 3.5 cm, and the intervals of time during which fish velocity is less than 1 BL/s are removed. Large leaps in fish trajectories during which fish move by more than 1.5 BL ≈ 5.25 cm between two consecutive frames, meaning that fish move at almost 65 cm/s, are also identified and removed, as they result from tracking errors. Finally, missing points are filled by linear interpolation. The final dataset used in this work represents approximately 10 hours of trajectory data for pairs of *H. rhodostomus*. For this work, trajectories of the original dataset are resampled with a timestep of Δ*t* = 0.12 s instead of the original 0.04 s provided by the camera, and data points have been converted from pixel space to a normalized [−1, 1] range to facilitate the training of the networks. The sub-sampling rate was chosen carefully to reduce the random noise between subsequent camera frames at the very short timescale of 0.04 s, while maintaining a sufficiently small timestep to study and model the social interactions.

### 4.2 Quantification of individual and collective behavior in pairs of fish

We use a set of observables to quantify how close the results of the models are from the measures obtained in the experiments [11, 19, 20]. These observables constitute a challenging benchmark when designing and testing a model. In the case of deep learning techniques, those observables also serve as means to partially explain what the algorithm has learned. In both cases, the observables constitute a stringent validation test.

Let us first define the temporal variables characterizing the individual and collective behavior of the fish. Fig. 4A shows two fish swimming in a circular tank of radius *R* = 25 cm. The position vector of a fish *i* at time *t* is given by its Cartesian coordinates 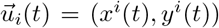 in the system of reference, centered at the center of the tank *C*(0, 0). The components of the velocity vector 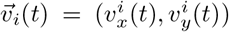 are given by 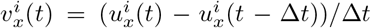 (similar expression for 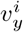). The heading angle of the fish is assumed to indicate its direction of motion and is therefore given by the angle that the velocity vector forms with the horizontal, 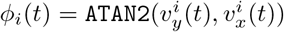.

**Fig. 4:**
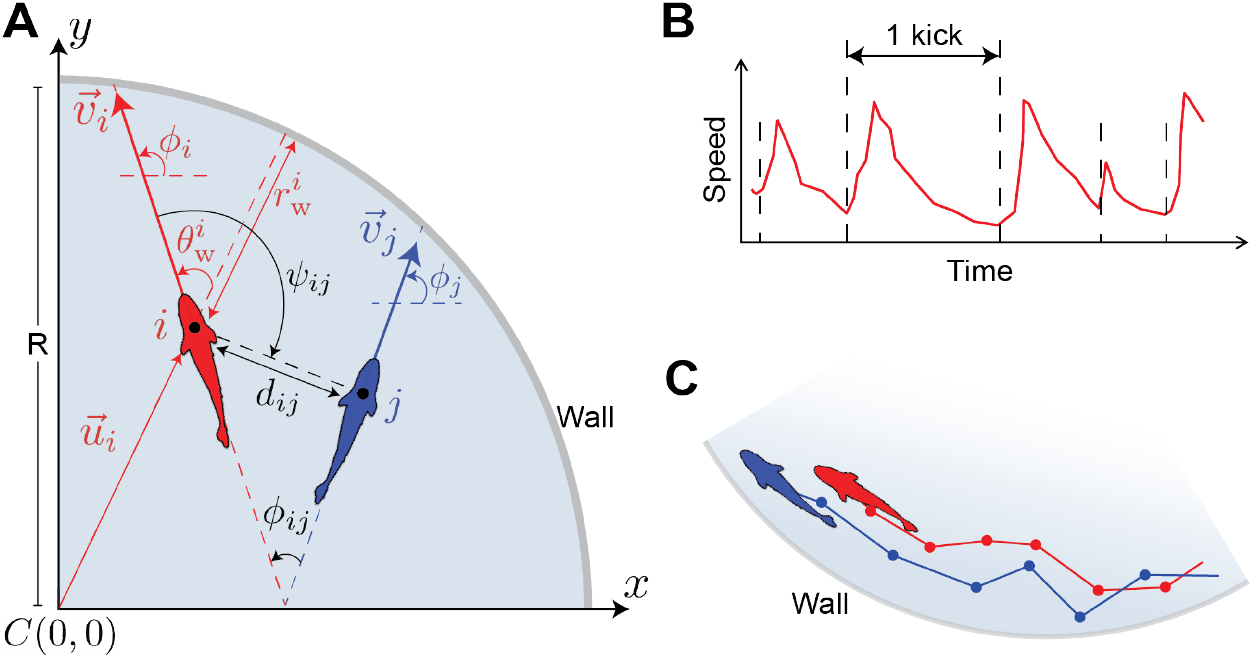
**A.** Individual and collective variables characterizing the instantaneous state of an individual (focal fish in red) and its pairwise relation with a neighbor (blue): distance to the wall 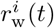, angle of incidence to the wall 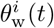, heading angle *ϕ_i_*(*t*), distance between individuals *d_ij_*(*t*), difference of heading angles *ϕ_ij_* (*t*), and angle of perception *ψ_ij_* (*t*). Positive angles (curved arrows) are defined in the anti-clockwise direction, starting from the positive semi-axis of abscissas. The radius of the circular setup is *R* = 25 cm. For visualization purposes, the size of fish is not to scale with the tank. **B.** Typical profile of the fish speed, *V*(*t*), showing the typical sequence of kicks (abrupt accelerations followed by longer gliding phases). **C.** Trajectories of two fish close to the wall due to their burst-and-coast swimming mode. The dots in the trajectories denote the instants of the kicks, where fish decision-making is assumed to take place.

The motion of a given fish *i* is then described using the three following instantaneous variables: the speed, 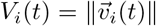, the distance of the fish to the wall, 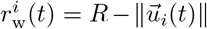, and the angle of incidence of the fish to the wall, 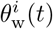, defined by the angle formed by the velocity vector and the normal to the wall: 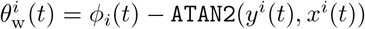, see Fig. 4A.

When there are two fish *i* and *j* in the tank, their relative motion is characterized by means of three variables: the distance between fish, 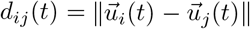, the difference between their heading angles, 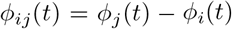, which measures the degree of alignment between both fish, and the angle of view, *ψ_ij_*(*t*), which is the angle with which fish *i* perceives fish *j*, and which is generally independent of *ψ_ij_*(*t*). See Fig. 4A for the graphical representation of these quantities. The angle of perception of the fish also allows us to define the notion of *geometrical leadership* for two fish: fish *i* is the *geometrical leader* (and therefore, *j* is the *geometrical follower*) if 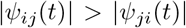, meaning that *i* has to turn by a larger angle to face *j* than the angle that *j* has to turn to face *i*. In practice, these definitions of the geometrical leader and follower provide a precise and intuitive characterization of a fish being ahead of the other. Note that being the leader or the follower is an instantaneous state that can change from one kick to the other.

These 6 quantities 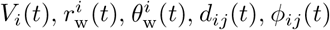, and 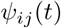 being defined, the measure of their probability distribution functions (PDF) constitutes a set of observables probing the individual and collective instantaneous fish dynamics in a fine-grained and precise manner. The PDF of 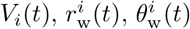 probe the behavior of a focal fish sampled over the observed dynamics, and are hence called *instantaneous individual observables*. The PDF of *d_ij_*(*t*), *ϕ_ij_*(*t*), and *ψ_ij_*(*t*) characterize the correlations between 2 fish *at the same time t* and are hence called *instantaneous collective observables*. These 3 collective observables can be easily generalized to a group of arbitrary size *N* > 2, by considering *i* and *j* as pairs of nearest neighbors, or pairs of second-nearest neighbors (or even farther neighbors), or even averaging them over all pairs in the group (then probing the size, the polarization, and the anisotropy of the group). Ultimately, comparing experimental results and model predictions for these individual and collective observables constitute a stringent test of a model.

Moreover, to characterize the *temporal correlations* arising in the dynamics, we make use of 3 additional observables involving quantities measured *at two different times*, for a given focal fish [20]: the mean-squared displacement *C_X_*(*t*), the velocity autocorrelation *C_V_*(*t*), and, especially challenging, the autocorrelation of the angle of incidence to the wall *C*_*θ*_w__ (*t*), defined respectively by

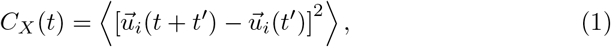

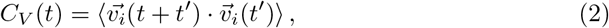

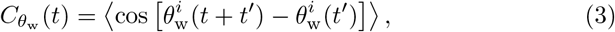

where 〈*w*(*t*)〉 is the average of a variable *w*(*t*) over all reference times *t*′ (assumption of a stationary dynamics, where correlations between two times depend solely on their time separation), over all focal fish, and over all experimental runs. Note that, although *C_V_*(*t*) can be shown to be the second derivative of *C_X_*(*t*), both quantities are measured independently. In principle, these correlation observables can also be generalized to probe the (collective) time correlations between the two different fish (or between nearest neighbors in a group of *N* > 2 individuals). For instance, one could consider 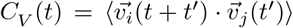, where the average is now over nearest neighbor pairs. However, in the present study, we will limit ourselves to the study of the 3 (individual) correlation functions listed in equations (1–3).

### 4.3 Analytical and deep learning models of fish behavior

Many species of fish like *H. rhodostomus* or *Danio rerio* move in a *burst-and-coast* manner, meaning that their swimming pattern consists of a sequence of abrupt accelerations each followed by a longer gliding period (Fig. 4B), during which a fish moves more or less in a straight line (Fig. 2C). The kicking instants observed in the curve of the speed can be interpreted as decision times when the fish potentially initiates a change of direction. In *H. rhodostomus*, the mean time interval between kicks and the typical kick length were experimentally found to be close to 0.5 s and 7 cm, respectively [11]. When confined in circular tanks, fish tend to swim close to the curved wall because their trajectory is made of quasi straight segments with limited variance of the heading angle between kicks, hence preventing the fish to escape from the tank wall (unless when a rare large heading angle change occurs) [11]. When swimming in groups, *H. rhodostomus* tend to remain close to each other, especially when the number of fish in the tank is small. In fact, the social interactions between fish reflect the combined tendency to align with and follow their neighbors while at the same time maintaining a safe distance with the wall. At a given kicking instant, only a few neighbors (one or two) have a relevant influence on the behavior of a fish [32]. The decision-making of fish displaying a burst-and-coast swimming mode can thus be reproduced by considering only pairwise interactions. Obviously, if one only considers pairs of fish, like here, it therefore suffices to consider the relative state of the neighboring fish (relative position and velocity) and the effect of the distance and the relative orientation to the wall [11, 19].

#### 4.3.1 Analytical Burst-and-Coast model

The Analytical Burst-and-Coast model (hereafter called ABC model) quantitatively reproduces the interaction dynamics of *H. rhodostomus* swimming alone or in pairs under the hypothesis that fish decision-making times correspond exactly to their kicking times, that is, the new direction of movement, the duration, and the length of the kick are decided precisely at the end of the previous kick [11].

Given a pair of agents *i* and *j* at a respective state 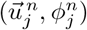 and 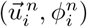 at time *t^n^*, the state of agent *i* at the next instant of time 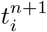 is given by

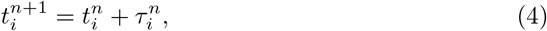

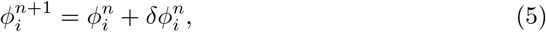

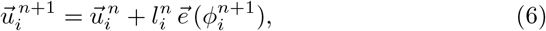

where 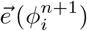 is the unitary vector pointing in the heading direction 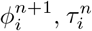 and 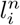 are the duration and length of the *n*-th of agent *i* kick, and 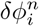 is the heading change of agent *i*. The heading angle change 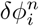 is the result of three effects: the effect of the wall, the effect induced by the social interactions with the other fish (repulsion/attraction and alignment), and the natural spontaneous fluctuations of fish motion (cognitive noise). The social influence depends only on the relative state of both agents, determined by the triplet 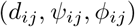. The derivation of the shape and intensity of the functions involved in 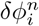 is based on physical principles of symmetry of angular functions and a data-driven reconstruction procedure detailed in [11] for the case of *H. rhodostomus* and in [19] for the general case of animal groups.

Starting from the initial condition 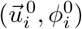 of fish *i*, the length and the duration of its next kick, 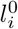 and 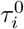, are sampled from the experimental distributions obtained in [11]. Then, the timeline 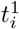 of fish *i* is updated with equation (4), the heading angle of the next kick 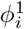 is calculated with equation (5), and the position of the fish at the end of the kick 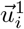 is obtained with equation (6). As kicks of different fish are asynchronous, the next kick can be performed by any of the two fish. Each fish has thus it own timeline, but is subject, at each of its kicks, to the evolution of the other fish along its own kicks.

The ABC model is therefore a discrete model that generates kick events instead of continuous time positions. To directly compare with the DLI model presented in the next section, which is a continuous time model, we re-sampled the trajectories made of kick events produced by the ABC model and build continuous time trajectories with a timestep of size Δ*t* = 0.12 s. We produced trajectories that add up to a total of approximately 16.6 hours of duration, that is, 500,000 timesteps.

#### 4.3.2 Deep Learning Interaction model

The Deep Learning Interaction model (hereafter called DLI model) consists of an Artificial Neural Network (ANN) which is fed with a set of variables characterizing the motion of *H. rhodostomus* and which provides the necessary information to reproduce the social interactions of these fish by estimating their motion along timestep of length Δ*t* = 0.12 *s*. At time *t*, the DLI is designed to take sequences of states as input to capture the short- and longterm dynamics. Then, it generates predictions for the acceleration components of the fish at the following timestep *t* + Δ*t*.

For the DLI model, the state of an agent *i* at time *t* is defined by

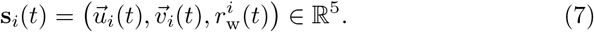

The state of an agent includes redundant information: in a fixed geometry, 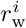 can be deduced from 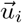, and 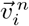 from the input sequence 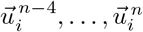. This redundancy is intended to facilitate the training process of the neural network.

The system’s state **S**(*t*) is then defined as the combination of both agent states, in addition to their inter-individual distance *d_ij_*(*t*) (also a redundant variable):

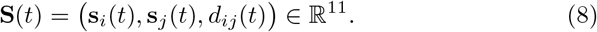

Fig. 5 shows the structure of the ANN, consisting of 7 layers: two Long-Short Term Memory (LSTM) layers [40], and 5 fully connected (Dense) layers. The first LSTM layer consists of 256 neurons and is located at the input of the ANN, where it receives the sequence of the 5 last states of the system, *i.e*., a matrix of dimension 5 × 11: (**S**(*t* – 4), …, **S**(*t*)). This history length of 4 timesteps (0.48 s) is borrowed from the biology of the fish: as already mentioned, the time it takes for a fish to display its characteristic behavior, a kick, is 0.5 s [11], therefore, we input the current state plus the states that correspond to the average duration of a kick. The output of the first LSTM is then gradually reduced in dimension by two successive dense layers, and then scaled up again with a second LSTM, whose configuration is also based on a history of 5 states. Then, two other dense layers are used to reduce the dimension of the output of the second LSTM, and a last dense layer is applied to provide the final output of the ANN. More details about the configuration of the ANN are given in Supplementary Table 1.

**Fig. 5:**
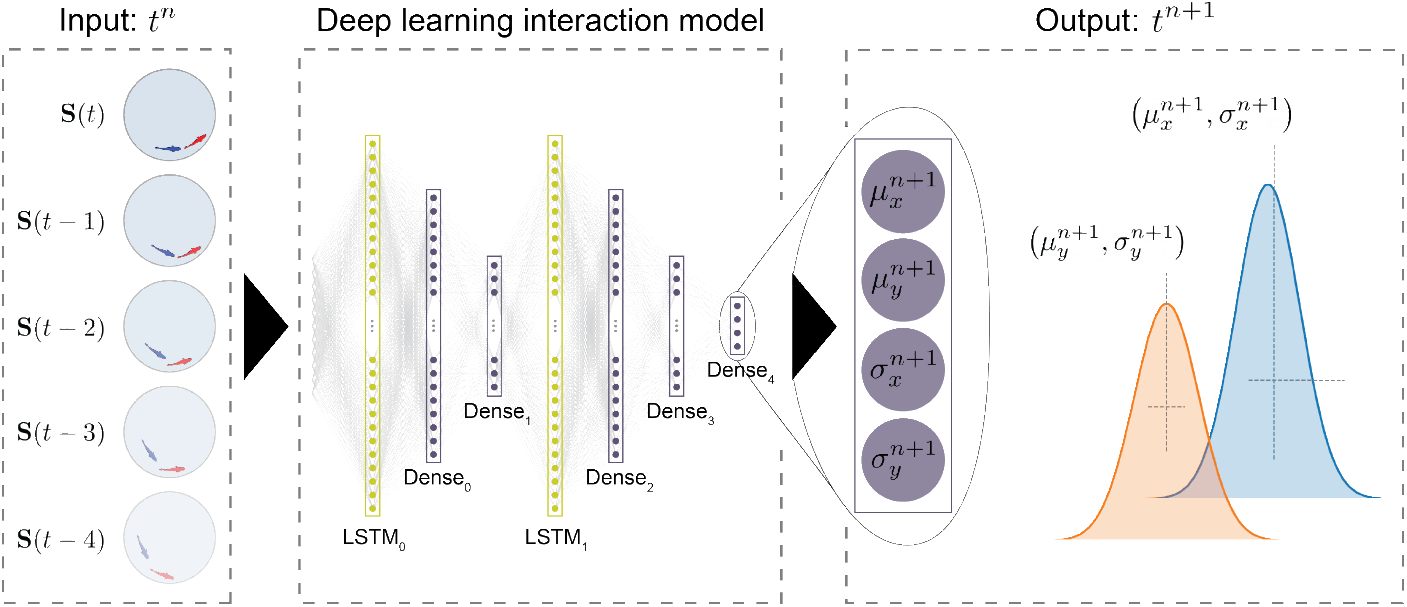
Structure of the Artificial Neural Network (ANN) used in the DLI model. From left to right: *Input* of the ANN: the 5 last states, (**S**(*t* – 4), …, **S**(*t*)) at time *t*. Where 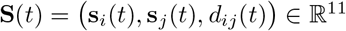 and each state is parametrized as 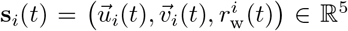; the 7 layers (two LSTM layers and 5 Dense Layers) capturing the social dynamics; *Output*: the two pairs of values (*μ_x_*, *σ_x_*) and (*μ_y_*, *σ_y_*) corresponding respectively to the mean and standard deviation of the probability distribution function (assumed to be Gaussian) of each component *a_x_* and *a_y_* of the instantaneous acceleration vector 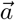 at time *t* + 1, constituting the prediction of the DLI model.

The output of the ANN consists of two pairs of values, (*μ_x_*, *σ_x_*) and (*μ_y_*, *σ_y_*), corresponding to the expected value and standard deviation of the *x* and *y* components of the predicted acceleration, which are assumed to be Gaussian distributed [41], as actually found for *H. rhodostomus* [11]. Hence, the predicted acceleration of the agent, 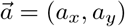, can be written

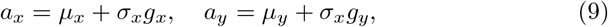

where *g_x_* and *g_y_* are independent standard Gaussian random variables drawn from 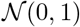. Then, the velocity vector of the agent *i* at the time *t*^*n*+1^ is given by

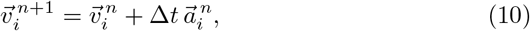

and the position of the agent is updated according to

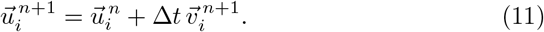

Note that in the DLI model, the predicted variance of the acceleration accounts for the fish intrinsic spontaneous behavior exhibited during their decision process (cognitive noise), and hence translates the fact that 2 real (or modeled) fish will not act the same if put twice in the same given state characterized by equation (8).

The *prediction* of the ANN for at time *t*^*n*+1^ is thus a vector of dimension 4 × 1 that can be written as 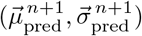, where

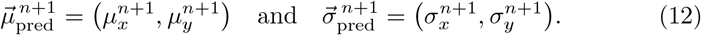

The ANN is then trained to approach the *real/observed* values 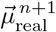 by means of the Adaptive Moment Estimation Optimizer (Adam) with a time-decaying learning rate λ = 10^−4^ and a negative log-likelihood loss function ℓ defined in terms of the prediction error 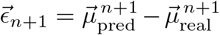 and the standard deviations as follows [42]:

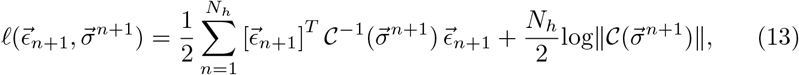

where *N_h_* is the number of timesteps in the history of the input of the ANN (here *N_h_* = 5) and 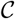 is a diagonal covariance matrix with the values of 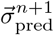 in the diagonal and zeroes elsewhere.

The training of the ANN is carried out with a subset of the experimental dataset. More specifically, the training process is given a budget of 45 epochs with a batch size of 512 samples on a dataset that was split 80%, 15%, and 5% for training, validation, and test, respectively. Then, the DLI is used to produce trajectories of 500,000 timesteps of size Δ*t* = 0.12 s, as done with the ABC model. At the beginning of the simulation, each agent is given a copy of the DLI model and both agents are initialized with a random 5 timesteps long trajectory. At each timestep *t^n^*, the state vector **S**(*t^n^*) is built and introduced in the network, which provides the estimated instantaneous acceleration distributions at time *t*^*n*+1^. Then, the acceleration is evaluated according to equation (9), and the next positions of the agents 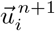 and 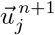 are obtained from the equations of motion, equations (10,11).

##### Designing the DLI model

Designing and selecting an appropriate ANN structure to model a system is for the most part non-trivial and requires either an extensive search through automatic methods (*e.g*., neuro-evolution [43–45]) or an exhaustive number of empirical attempts for very specific applications [21–23]. Here, we followed a hybrid approach consisting in empirically designing an ANN based on biological insight and automatically searching for its optimal structure by bootstrapping the search. Once we established this initial model, we performed an automated search for similar neural networks using the same input and output for different combinations of *i*) the number of layers, *ii*) the size of the layers, and *iii*) the activation functions (*i.e*., transfer functions tasked with mapping the inputs of a neuron to a single weighted output value passed to the next layer). The search included a total of 82 neural network structures, out of which the ANN shown above is the best performing.

Three notable categories of networks were considered: *i*) non-probabilistic networks that only generate 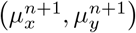 (and hence, not explicitly including the cognitive noise), *ii*) probabilistic networks that do not have memory cells (hence, missing the fact that fish are gliding passively on a timescale of order 0.5 s), and *iii*) probabilistic networks that implement memory thanks to LSTM layers. Non-probabilistic networks (*i*) provide the mean value of the components of the acceleration for the next timestep with high accuracy, but miss the essential variability that is intrinsic to the spontaneous behavior of fish and which allows for the emergence of social interactions. Probabilistic networks without memory (*ii*) are able to partly capture this intrinsic variability, but do not fully capture the non-linear nature of the problem (see Supplementary Figure 4 and Supplementary Video 3). Finally, probabilistic networks with memory (*iii*) performed generally well, and we found that the structure used in the DLI model consistently provides the best results for the number of epochs set for training and for the ANNs considered by the automatic search.

Our search approach revealed the existence of two crucial ingredients that must be considered in the model, both accounting for biological characteristics of fish behavior observed experimentally. First, the neural network must be fed with information covering the typical timescale along which relevant changes take place in the behavior of the fish. Since real fish kicks last 0.5 s on average, the NN needs information about the fish behavior over time intervals of at least this duration (that is, 4 to 5 timesteps of 0.12 s). However, we found that using longer vector lengths (up to 10 timesteps) for the case of *H. rhodostomus* does not lead to any significant improvement in the results, while considerably increasing the training time. Second, the output of the network must contain a sufficiently wide diversity of predictions so that the agents reproduce the high variability of responses that fish display when behaving spontaneously and reacting to external stimuli.

ANNs without memory tend to make too similar predictions, and agents do not initiate the typical direction changes that are observed in the experiments. A possible solution could be to add some phenomenological noise to the predictions of the NN. However, this would result in an unrealistic behavior, albeit an improvement over not adding noise at all. For example, when a fish swims close to the wall, it does not have the same liberty to turn toward or away from the wall, which would not be captured by a too crude implementation of the fish cognitive noise. Our approach accounts for this behavioral uncertainty for each state (position, velocity, distance to the neighbor and to the wall) and for both degrees of freedom during the training phase of the ANN, being therefore able to capture these complex behavioral patterns. The performance of the two variants is depicting in Supplementary Figure 4.

## Ethics approval

The experiments conducted with *H. rhodostomus* were approved by the Ethics Committee for Animal Experimentation of the Toulouse Research Federation in Biology no. 1 and comply with the European legislation for animal welfare. The experiments conducted with *D. rerio* were approved by the state ethical board of the Department of Consumer and Veterinary Affairs of the Canton de Vaud (SCAV) of Switzerland (authorization no. 2778).

## Code & data availability

All the code concerning the data pre-processing, neural networks, and plot scripts is publicly available in https://doi.org/10.5281/zenodo.7634914. Experimental and generated data are available in https://doi.org/10.5281/zenodo.7634688.

## Acknowledgments

We would like to thank Dr. Frank Bonnet for his contribution in the early stages of this paper and the Franco-Swiss collaboration between the École polytechnique fédérale de Lausanne (V.P. and F.M.), the Centre de Recherches sur la Cognition Animale (R.E. and G.T.) and the Laboratoire de Physique Théorique (C.S.) at Université Paul Sabatier.

## Authors’ contributions

### Study concept & design

F.M., V.P., C.S., G.T conceived and designed the study.

### Code implementation & analysis

V.P. implemented the pre-processing, neural networks, plot scripts and concept figures. C.S. also assisted with the plot scripts and implemented the simulation of the model from [11]. The design and fine-tuning of the neural network structures was done by V.P. A.A. and V.P. configured the D-LSTM. V.P. and C.S. analyzed the data.

### Experiments

C.S., G.T. conducted the experiments and their analysis with *H.rhodostomus*. V.P. conducted the experiments with *D. rerio*.

### Writing - Original draft

V.P., C.S.

### Writing - Review & editing

A.A., R.E., F.M., V.P., C.S., G.T.

## Supplementary information

### 1 Supplementary Videos

**Supplementary Video 1:** Examples of trajectories obtained in experiments with *H. rhodostomus* (left), for the Analytical burst-and-coast (ABC) model (center), and for the Deep Learning Interaction (DLI) model (right). This video illustrates the qualitative agreement between trajectories generated by the ABC and DLI models and experimental trajectories, while the quantitative agreement between the models and experiments is studied in detail in the Result section. The video can be downloaded at https://github.com/epfl-mobots/ncs_preddl_2023/raw/main/Videos/SI_Video1.mp4.

**Supplementary Video 2:** Example of a generated trajectory simulation for the D-LSTM model. Already at the qualitative level, the D-LSTM model fails at reproducing realistic trajectories (compare with Supplementary Video 1). The video can be downloaded at https://github.com/epfl-mobots/ncs_preddl_2023/raw/main/Videos/SI_Video2.mp4.

**Supplementary Video 3:** Example of a generated trajectory simulation for the Multi-layered Perceptron Interaction (MLI) model. Already at the qualitative level, the MLI model fails at reproducing realistic trajectories (compare with Supplementary Video 1). The video can be downloaded at https://github.com/epfl-mobots/ncs_preddl_2023/raw/main/Videos/SI_Video3.mp4.

**Supplementary Video 4:** Example of collective behavior in a group of 5 DLI agents, *without any retraining*. For a given focal agent, we compute the predicted acceleration and noise which would be produced by each of the 4 other agents. Following [32], we define the two most influential neighbors as the neighbors leading to the two highest predicted accelerations. Ultimately, the focal fish speed and position are updated according to equations (9–11), using the sum of these two highest accelerations and the average predicted noise. This video illustrates the fact that, although the DLI was only trained to mimic the social interactions between pairs of fish, it produces cohesive and aligned trajectories for 5 agents, in good qualitative agreement with corresponding trajectories for 5 rummy-nose tetra [32]. In the future, we will address the quantitative comparison between long-term trajectories for groups of DLI agents and real fish, in particular, in connection to our robotic platform [34–36]. The video can be downloaded at https://github.com/epfl-mobots/ncs_preddl_2023/raw/main/Videos/SI_Video4.mp4.

### 2 Supplementary Notes 1

Along with ABC, we also adapted a neural network from [30], that was initially intended for human trajectory forecasting in crowded spaces, for pair-wise fish interactions. More specifically, we used a D-MLP-ConC-LSTM as described in [30], which we refer to as D-LSTM in short. The models presented in [30] were introduced in the context of forecasting human trajectories in arbitrary scenes of humans walking. We found that there are many similarities in the approach and goals of human trajectory forecasting research works and the goals of our work, *i.e*., to model the interactions between fish. Therefore, we chose to adapt and use this algorithm to obtain a baseline of the performance that our neural network, presented in the following subsection, achieves. We opted to not intervene with the core structure of the neural network and use the framework provided by the authors of [29, 30]^1^, and parametrize it to generate accurate trajectory estimates. An example of generate trajectory simulation for the MLI can be found in Supplementary Video 2.

The model’s name is partly owed to its directional pooling layer, where the relative velocity of neighboring individuals is combined with the focal individual’s to estimate its future trajectory. To obtain predictions, we trained the network with the same input data, only adapted to a different format compatible with the TrajNet++ framework [30]. Furthermore, we opted to convert the inputs to meters, instead of the arbitrary [−1, 1] scale, to allow for more intuitive parameter selection. The TrajNet++ framework provides a series of different tools for different tasks, including pre-processing of datasets. Before training the D-LSTM, we performed a pre-processing methodology provided by TrajNet++ that categorizes the data in 4 categories depending on the trajectory type. Namely, static (Type I), linear (Type II), non-linear Type III) and non-interacting (Type IV) trajectories. Here, we parametrized the percentage of the trajectories that correspond to those types and are subsequently used for training, with values 80%, 100%, 100%, and 100%, respectively. Then, the TrajNet++ tool split the dataset in 80-15-5% manner for the training, validation and test datasets. We also set the frame rate option for the pre-processing tool to 25 frames per second, which corresponds to the timescale chosen for ABC, DLI, and the experiments. Finally, we parametrized the group distance threshold to 0.5 m, that is, the diameter of the setup to always consider both individuals in the dataset.

Then, similarly to DLI, the parameters of the model were learned by minimizing a negative log-likelihood loss using the Adam optimizer [42] with an initial learning rate of λ = 0.0001. A step learning rate scheduler was used to reduce the learning rate with every 10 epochs. The network was given a budget of 80 epochs and a batch size of 8. With respect to the directional pooling layer and the underlying grid, we selected a grid of 32 cells with a cell side size of 0.005 m (see details in [30]).

The network itself is structured as follows; 1) the input state, rectangular position, of the focal individual is passed to a Multi-layered perceptron (MLP) structure. We opted to maintain the original input information of the D-LSTM, therefore, contrary to the DLI, the D-LSTM is missing 3 input variables, namely, distance to the wall for the focal and the neighboring individual, and inter-individual distance. 2) the directional information (relative velocity and position) of the top-k (in the context of this work *k* = 1) neighbors are concatenated, and 3) the concatenation is passed to an LSTM layer. Both the MLP and LSTM layers consist of 256 neurons. The network was given observations of length 5, similarly to DLI, and asked to predict the mean and standard deviation of the components of acceleration of 3 future time steps (duration of Δ*t* = 0.36 s), instead of 1 that was required from DLI, for two reasons; 1) to put pressure during training on the network to learn more accurately, and 2) to compare its longer horizon forecasting capabilities against DLI. In D-LSTM’s simulations, we considered only the prediction corresponding to *t* + Δ*t* and followed the same simulation logic of DLI (see Section 4.3.2). That means, a new trajectory was generated at every time step (Δ*t* = 0.12 s). It is indeed common, in such cases, that only one time step is used for simulation and the remaining part of the trajectory is used to understand the estimated intent of the agent.

#### 2.1 Comparing the short- and long-term performance of DLI and D-LSTM

##### 2.1.1 Quantification of instantaneous individual behavior

D-LSTM does not very well produce agents that swim at the same speed as fish in the experiments. The location of the peak of the PDF of V (see Supplementary Figure 1A) is indeed in good agreement with the experiments, but the distribution decays much faster than in the experiments and DLI and goes to zero at approximately 25 *cm/s*. Supplementary Figure 1B shows that the PDFs of the D-LSTM leader and follower are not in good agreement with the experiments or DLI and both agents swim very close to the wall most of the time. Furthermore, agents tend to swim at a farther distance from the wall more often than in the experiments (see the dashed and dotted lines in Supplementary Figure 1B between 6 – 25 *cm*) and the DLI. Similarly, Supplementary Figure 1C shows that alignment with the wall is not well reproduced by the D-LSTM model. The locations of the peaks are well reproduced with respect to the experiments, including an angle of incidence *θ*_w_ smaller than 90°. However, the D-LSTM agents swim parallel to the wall considerably less than in the experiments, see the PDF for values of *θ*_w_ < −90° and *θ*_w_ > 90°.

##### 2.1.2 Quantification of instantaneous collective behavior

The PDF of the inter-individual distance *d_ij_* is quantitatively different from what is measured for the experiments. Notably, the location of the peak is located approximately 1, *cm* more to the left (i.e., closer to the neighboring agent) than in the experiments (see Supplementary Figure 1). Furthermore, the D-LSTM agents swim much closer to each other and almost never swim more than 15 cm away from each other, contrary to the experiments. Conversely, the D-LSTM reproduces the alignment PDF *ϕ_ij_* very well, albeit D-LSTM agents tend to be unaligned more often than in the experiments and DLI. Similarly, to ABC and DLI, the D-LSTM fails to reproduce the small increase in the PDF at Δ*phi* ≈ 180°. Viewing angles are very well reproduced by the D-LSTM. In fact, the PDF is very similar to those of ABC and DLI, although the D-LSTM follower agent (see dotted lines in Supplementary Figure 1) tends to swim less often parallel to the leader agent.

**Supplementary Figure 1:**
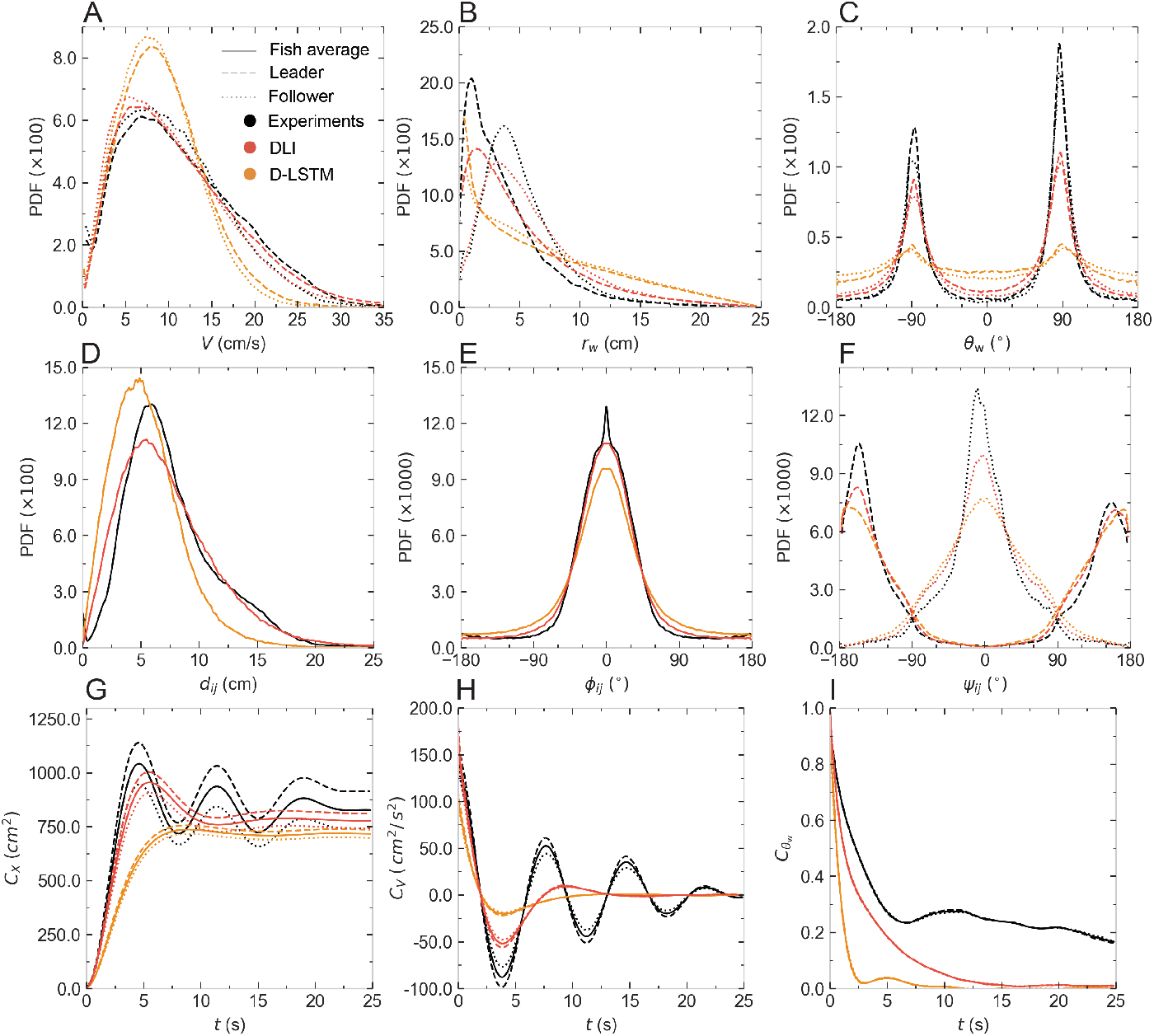
Probability density functions (PDF) of all observables. **A** Speed *V*, **B** distance to the wall *r*_w_, **C** angle of incidence to the wall *θ*_w_, **D** Distance between individuals *d*, **E** difference in heading angles *ϕ_ij_*, **F** angle of perception of the geometrical leader and follower *ψ*, **G** Mean squared displacement (*i.e*., 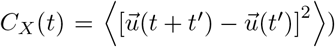, **H** Velocity temporal correlations (*i.e*., 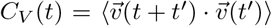), and **I** Temporal correlations of the heading of a fish relative to the wall (*i.e*., 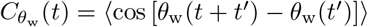). Black lines: experimental data of real fish. Orange lines: agents of D-LSTM. Red lines: agents of DLI (Deep learning interaction model). Dashed lines correspond to the leader, and dotted lines to the follower.

##### 2.1.3 Quantification of time correlation

Supplementary Figure 1G shows that the D-LSTM cannot well reproduce the short or intermediate regimes of *C_X_*(*t*). The curve saturates later than the fish in the experiments (2 s more) being considerably farther distance from the wall compared to fish or ABC and DLI. Similarly to fish, ABC and DLI, the leader and follower have different curves for *C_X_*(*t*) and *C_V_*(*t*) (Supplementary Figure 1G and H, respectively), although in D-LSTM the effect is less pronounced than the other two interaction models. The oscillations of *C_V_*(*t*) in D-LSTM are sharply damped and disappear at *t* = 5 s. Similarly to ABC and DLI, the oscillations in *C*_*θ*_w__ (*t*) are not in good agreement with fish data and fail to reproduce the correlation of fish for *t* > 2.5 *s*.

#### 2.2 Comparing the short-term performance of DLI and D-LSTM

Trajectory forecasting algorithms provide a longer trajectory horizon than *t* + Δ*t* in an attempt to capture the intent of the agents. Here, the ANNs’ goal is to reproduce similar movement trajectories as fish.

In Supplementary Figure 2, we depict a box-plot of the performance of DLI and D-LSTM with respect to their mean squared error measured against the fish in the experiments. Both models have achieved comparable performance as the D-LSTM performing systematically better by marginal difference. As expected, the models demonstrated an increasing error (see Supplementary Figure 2A) for the more distant horizon (i.e, at 0.36 *s*). We also depict a few sample instances of both models in action in Supplementary Figure 2B(i-vi), along with their uncertainty maps (on the right-hand side of each example). Supplementary Figure 2A show that D-LSTM make more confident decisions, that is, the generated trajectories exhibit less variability. After visual qualitative inspection of approximately 3.500 trajectory samples, such as the ones of Supplementary Figure 2B, we did not observe any differences or particularities for either model.

**Supplementary Figure 2:**
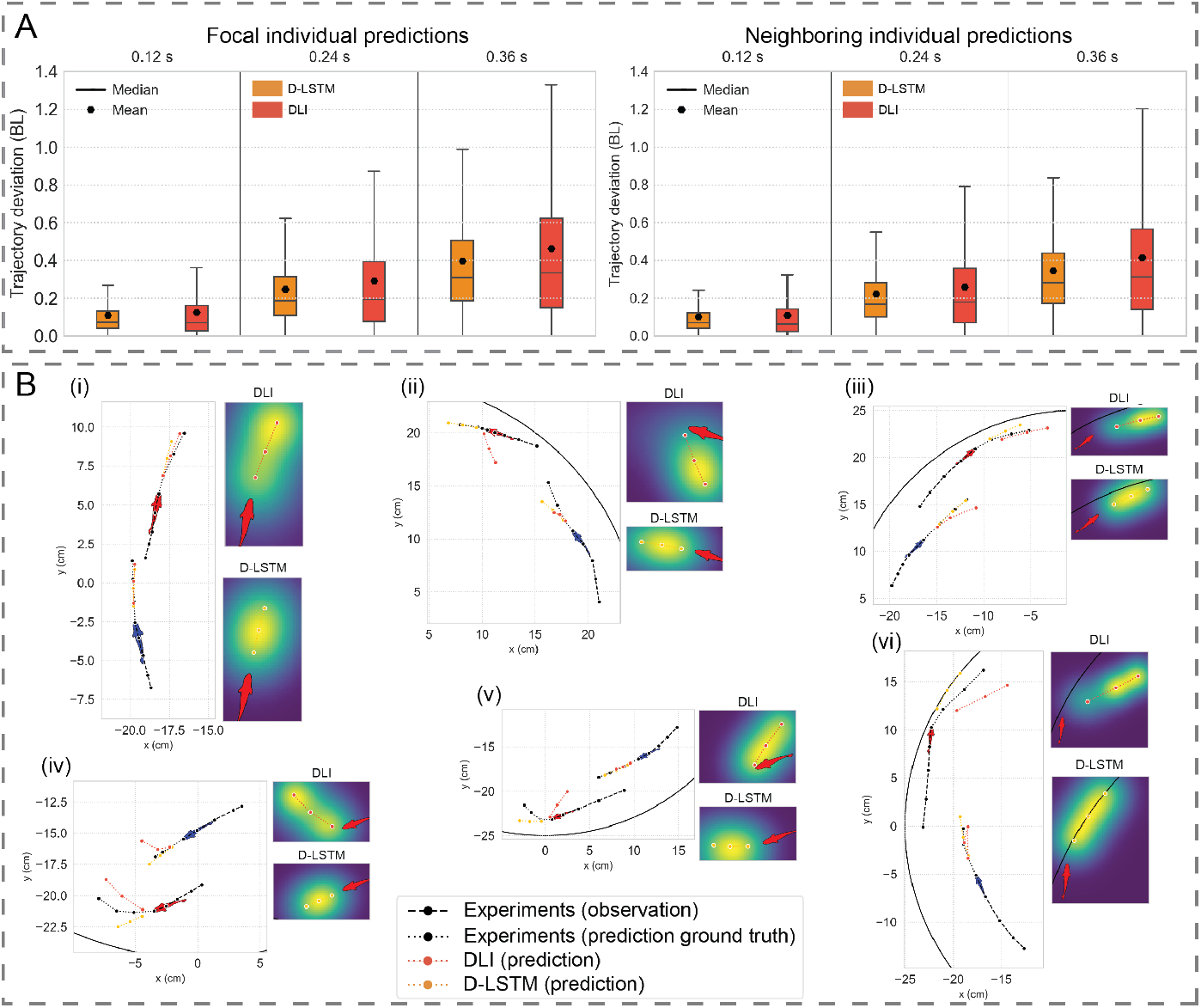
**A** Mean squared error between the generated position and the actual position of the focal individual (left) and the neighbor (right), and **B** examples of generated trajectories for DLI and D-LSTM (left) and uncertainty of the models (right) for all panels **(i)-(vi)**, where certainty is depicted with hues of blue to yellow, for low to high certainty areas, respectively.

### 3 Supplementary Notes 2

The premise of ML algorithms is that they can more easily scale up to solve similar tasks. In this subsection, we put this to the test. That is, to validate the DLI’s performance with a different fish species. Therefore, we conducted experiments of *D. rerio* pairs, similarly to the *H. rhodostomus*. Then, we trained the DLI as described in Section 4.3.2.

**Supplementary Figure 3:**
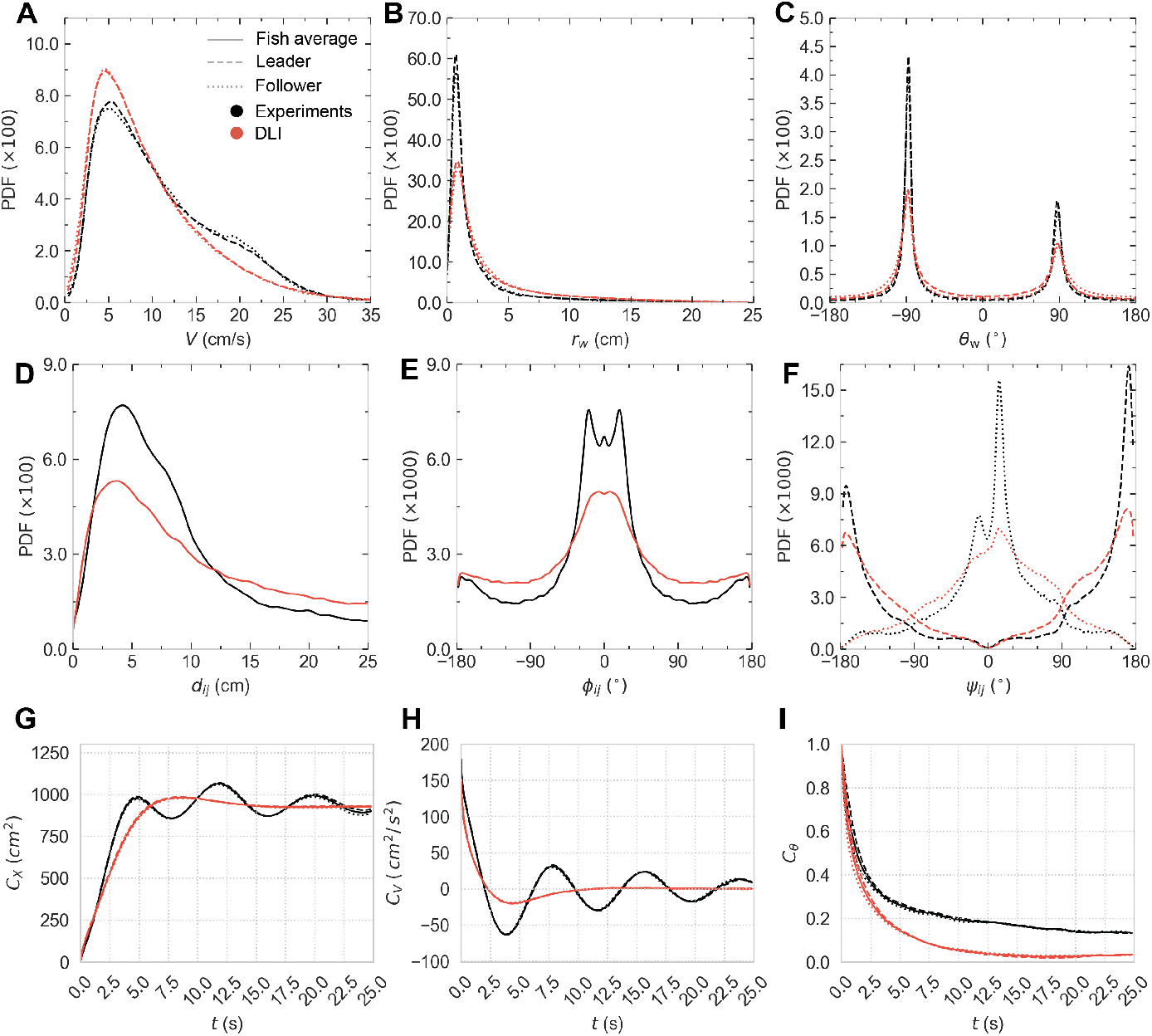
Probability density functions (PDF) of all observables. **A** Speed *V*, **B** distance to the wall *r*_w_, **C** angle of incidence to the wall *θ*_w_, **D** Distance between individuals *d*, **E** difference in heading angles *ϕ_ij_*, **F** angle of perception of the geometrical leader and follower *ψ*, **G** Mean squared displacement (*i.e*., 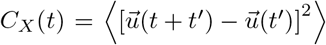), **H** Velocity temporal correlations (*i.e*., 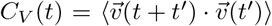), and **I** Temporal correlations of the heading of a fish relative to the wall (*i.e*., 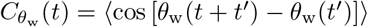). Black lines: experimental data for zebrafish. Red lines: agents of DLI. Dashed lines correspond to the leader, and dotted lines to the follower.

#### 3.1 Quantification of instantaneous individual behavior

Supplementary Figure 3A is very well reproduced by the DLI, capturing the peak of the PDF correctly. Marginal difference is noticed in velocities between 15 and 30 cm/s. The DLI is also reproducing well the PDF of the leader and follower distance to the wall *r*_w_ (Supplementary Figure 3B), with marginal differences producing a wider PDF for the DLI. In Supplementary Figure 3C, the PDFs of the angle of incidence to the wall *θ*_w_ show good agreement, with marginal differences between angles of −90° and 90°.

#### 3.2 Quantification of instantaneous collective behavior

In terms of the collective behavior, the DLI is marginally worse when compared to real fish for the interindividual distance *d_ij_* of agents (see Supplementary Figure 3D). Similarly, the DLI is not fully capable of recovering the PDF heading angles exhibiting more occurrences in the tails of the PDF (*ϕ_ij_* < −50° and *ϕ_ij_* > 50°). However, when DLI leader and follower agents swim, they are able to reproduce the PDF of viewing angle *ψ_ij_* with good agreement to the original fish experiments.

#### 3.3 Quantification of time correlation

Contrary to *H. rhodostomus*, *D. rerio* data present only 2 distinct regimes: quasi-ballistic regime at short timescale (*t* ≲ 2 s) where 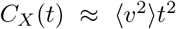, followed by a saturation regime (*t* > 5 s) characterized by slowly damped oscillations (Supplementary Figure 3G). Similarly, the velocity correlation function starts from 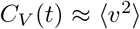 at short time and presents oscillations (Supplementary Figure 3H). Oscillations are eventually damped at a time greater than *t* ≫ 20 s. Contrary to *H. rhodostomus*, leader and follower showcase approximately the same saturation values. DLI agents reproduce the reproduce well the oscillations of *C_X_*(*t*) up to *t* ≈ 2.5 s. Contrary to the real experiments, the DLI PDF is quickly damped after this time. The same is true for *C_V_*(*t*), that is also quickly damped after *t* ≈ 2.5 s. However, the *C*_*θ*_w__ (*t*) PDF of the DLI model is in better agreement with the experiments, but the two curves deviate in amplitude at *t* ≈ 2 *s*

### 4 Supplementary Notes 3

One of the main alternatives investigated during the automated search described in Section 4.3.2 is a Multi-layered Perceptron Interaction (MLI) version of the DLI. That is, we maintained the general shape of the structure of 7 layers, but all of them are dense layers. That also means that the input is not a sequence of states, but only the last state at time *t^n^*. For reference, we also tested various architectures where a sequence of states was concatenated and provided at the input of the network, but we present MLI for two reasons: 1) concatenating past states at the input would provide some sort of memory to the ANN which is not the comparison we set out to make, and 2) the ANNs with sequence of states at the input (and no memory) did not give better results than MLI. Naturally, the lack of LSTM layers that consist of memory cells, means that the MLI is composed of significantly less free parameters than the DLI (approximately 1/6 th). An example of generate trajectory simulation for the MLI can be found in Supplementary Video 3.

#### 4.1 Quantification of instantaneous individual behavior

Supplementary Figure 4A shows that the PDF of V is considerably different than the experiments. More specifically, the location of the peak is located at a much lower velocity (≈ 1 *cm/s*). Similarly, the PDF of the MLI leader and follower of are not peaked at the same location as the experiments. However, the PDF of the MLI follower is peaked at a greater distance to the wall than the leader MLI agent, in agreement with what is measured in the experiments. Despite this, the PDF of the leader and follower also demonstrate smaller peaks around a distance of *d*_w_ = 19 *cm*. The alignment with the wall is also not very well reproduced by MLI (Supplementary Figure 4), although the asymmetry in the direction of rotation is captured to a small extent. Furthermore, whereas the PDFs of both the leader and follower MLI agent capture the slightly smaller angle of incidence *θ*_w_ when *θ*_w_ > 0, the angle of incidence for *θ*_w_ < 0 is approximately equal to 90°.

#### 4.2 Quantification of instantaneous collective behavior

In Supplementary Figure 4D the inter-individual distance *d_ij_* PDFs of MLI are not in good agreement with the experiments. Although the location of the peak is located at the same value as in the experiments, there is a second peak at *d_ij_* ≈ 26 *cm* which is not present in fish data. Similarly, Supplementary Figure 4E shows that MLI fails in reproducing the PDF of alignment *ϕ_ij_*, although the peak location is in good agreement with the experiments. However, MLI agents swim considerably less amount of time aligned than the experiments. The viewing angle (Supplementary Figure 4F) leader and follower PDF peaks are in good agreement with the experiments, but the asymmetry present in the fish data is not captured. Furthermore, both PDFs of the MLI are wider around the peaks than the experiments.

**Supplementary Figure 4:**
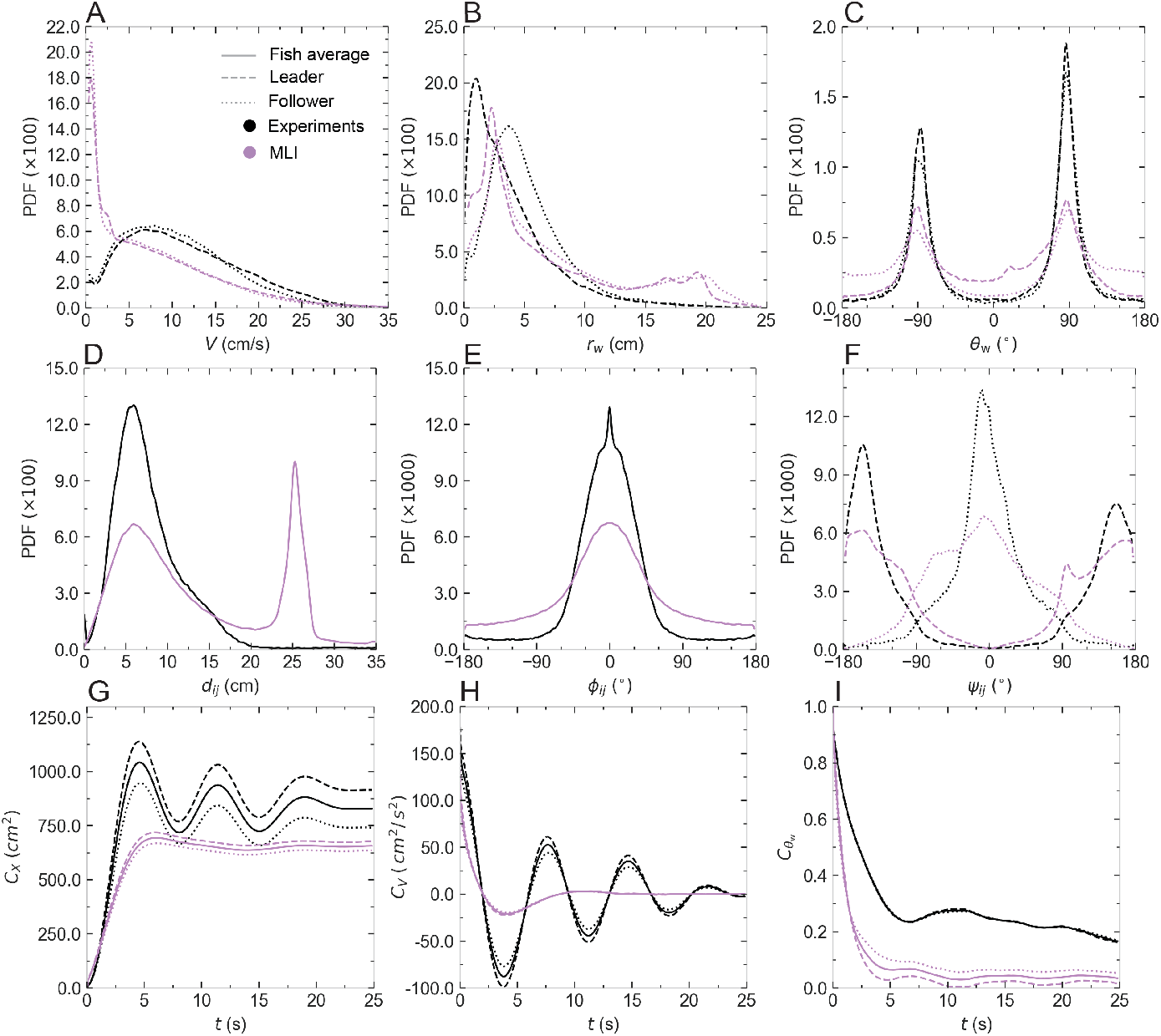
Probability density functions (PDF) of all observables. **A** Speed *V*, **B** distance to the wall *r*_w_, **C** angle of incidence to the wall *θ*_w_, **D** Distance between individuals *d*, **E** difference in heading angles *ϕ_ij_*, **F** angle of perception of the geometrical leader and follower *ψ*, **G** Mean squared displacement (*i.e*., 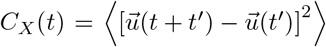), **H** Velocity temporal correlations (*i.e*., 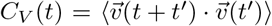), and **I** Temporal correlations of the heading of a fish relative to the wall (*i.e*., 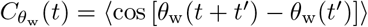). Black lines: experimental data of real fish. Orange lines: agents of D-LSTM. Red lines: agents of MLI (Multi-layered Perceptron Interaction model). Dashed lines correspond to the leader, and dotted lines to the follower.

#### 4.3 Quantification of time correlation

The MLI is not able to accurately reproduce the oscillations of *C_X_*(*t*) similarly to experiments. It saturates approximately 1 s later than the fish in the experiments, while considerably farther distance from the wall. Similarly to fish, ABC and DLI, the MLI leader and follower have different curves for *C_X_*(*t*) and *C_V_*(*t*) (Supplementary Figure 4G and H, respectively). Supplementary Figure 4H shows that MLI is also unable to reproduce the oscillations of *C_V_*(*t*). The curve deviates from *t* = 0 and reaches the first oscillation at time *t* = 5 *s*, similarly to the experiments, but is quickly damped after. *C*_*θ*_w__ (*t*) are not in good agreement with fish data and fail to reproduce the correlation of fish for *t* > 2.5 *s*. Notably, the leader and follower curves of the MLI are considerably different, in contrary to the fish data.

**Supplementary Table 1:**
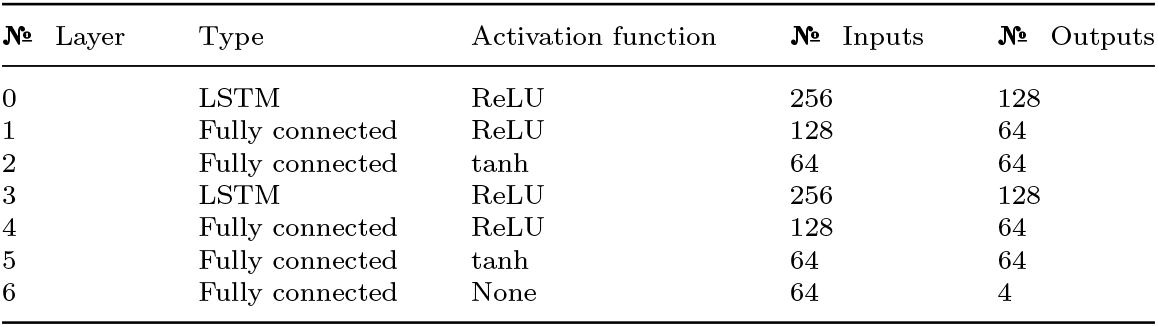
Implementation details of the DLI per layer. The 5 columns correspond to the increasing layer count, the type of layer, activation function, number of inputs, and number of outputs respectively.

**Supplementary Table 2:** Mean and standard deviation values for the experiment

| Quantity | Experimental dataset |  |  |
| --- | --- | --- | --- |
|  | Pair | Leader | Follower |
| **V (cm/s)** | $11.04 \pm 6.69$ | $11.54 \pm 7.05$ | $10.53 \pm 6.27$ |
| **r (cm)** | $4.58 \pm 3.53$ | $3.93 \pm 3.56$ | $5.23 \pm 3.38$ |
| $\theta_w$ (°) | $12.44 \pm 91.40$ | $12.07 \pm 90.25$ | $12.81 \pm 92.54$ |
| **d (cm)** | $8.16 \pm 5.76$ | — | — |
| $\phi_{ij}$ (°) | $0 \pm 53.47$ | — | — |
| $\psi$ (°) | $-8.54 \pm 107.47$ | $-15.66 \pm 143.82$ | $-1.42 \pm 48.12$ |

**Supplementary Table 3:** Mean and standard deviation values for the ABC model

| Quantity | ABC model dataset |  |  |
| --- | --- | --- | --- |
|  | Pair | Leader | Follower |
| **V (cm/s)** | $11.42 \pm 6.13$ | $11.62 \pm 6.18$ | $11.21 \pm 6.08$ |
| **r (cm)** | $4.93 \pm 4.20$ | $4.46 \pm 4.18$ | $5.40 \pm 4.17$ |
| $\theta_w$ (°) | $19.92 \pm 90.12$ | $19.48 \pm 88.02$ | $20.37 \pm 92.18$ |
| **d (cm)** | $7.92 \pm 5.44$ | — | — |
| $\phi_{ij}$ (°) | $0 \pm 40.72$ | — | — |
| $\psi$ (°) | $-5.45 \pm 107.41$ | $-9.87 \pm 144.03$ | $-1.02 \pm 47.85$ |

**Supplementary Table 4:**
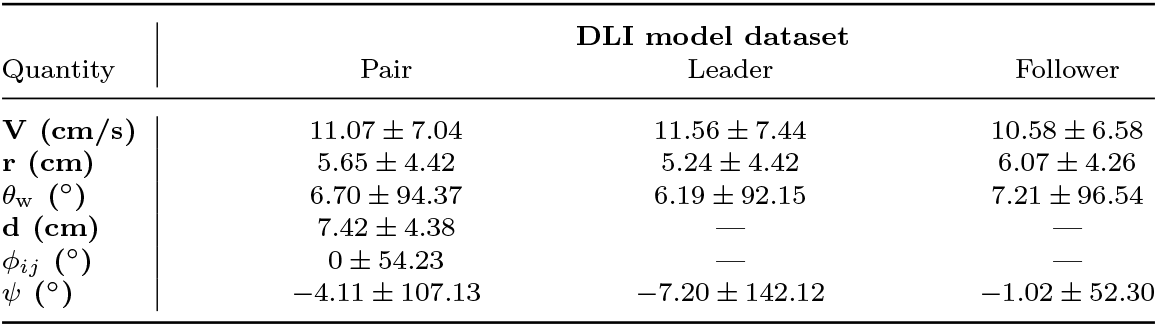
Mean and standard deviation values for the DLI model

| Quantity | DLI model dataset |  |  |
| --- | --- | --- | --- |
|  | Pair | Leader | Follower |
| **V (cm/s)** | $11.07 \pm 7.04$ | $11.56 \pm 7.44$ | $10.58 \pm 6.58$ |
| **r (cm)** | $5.65 \pm 4.42$ | $5.24 \pm 4.42$ | $6.07 \pm 4.26$ |
| $\theta_w$ ($^\circ$) | $6.70 \pm 94.37$ | $6.19 \pm 92.15$ | $7.21 \pm 96.54$ |
| **d (cm)** | $7.42 \pm 4.38$ | — | — |
| $\phi_{ij}$ ($^\circ$) | $0 \pm 54.23$ | — | — |
| $\psi$ ($^\circ$) | $-4.11 \pm 107.13$ | $-7.20 \pm 142.12$ | $-1.02 \pm 52.30$ |

**Supplementary Table 5:** Mean and standard deviation values for the Euclidean distance between prediction and real trajectory (focal individual)

| Future time-point | Focal individual |  |  |
| --- | --- | --- | --- |
|  | 0.12 s | 0.24 s | 0.36 s |
| DLI | $0.12 \pm 0.11$ | $0.29 \pm 0.26$ | $0.46 \pm 0.46$ |
| **D-LSTM** | $0.11 \pm 0.13$ | $0.25 \pm 0.22$ | $0.40 \pm 0.32$ |

**Supplementary Table 6:** Mean and standard deviation values for the Euclidean distance between prediction and real trajectory (neighboring individual)

| Future time-point | Focal individual |  |  |
| --- | --- | --- | --- |
|  | 0.12 s | 0.24 s | 0.36 s |
| DLI | $0.11 \pm 0.15$ | $0.26 \pm 0.27$ | $0.41 \pm 0.39$ |
| **D-LSTM** | $0.10 \pm 0.12$ | $0.22 \pm 0.20$ | $0.35 \pm 0.39$ |

**Supplementary Table 7:**
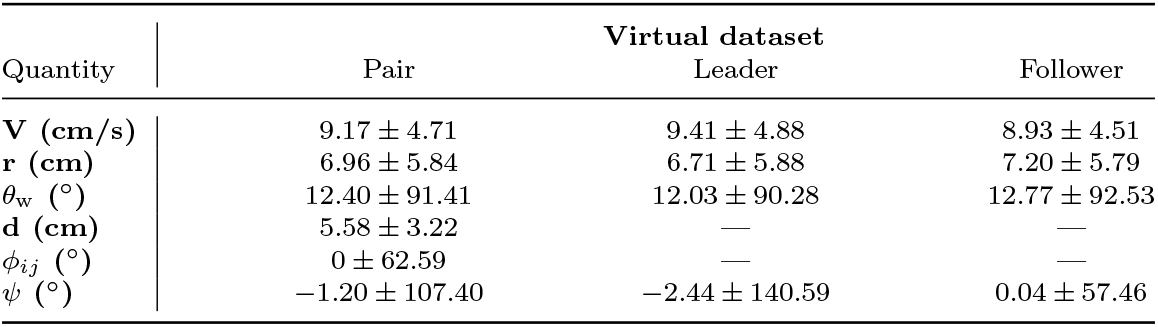
Mean and standard deviation values for D-LSTM

**Supplementary Table 8:**
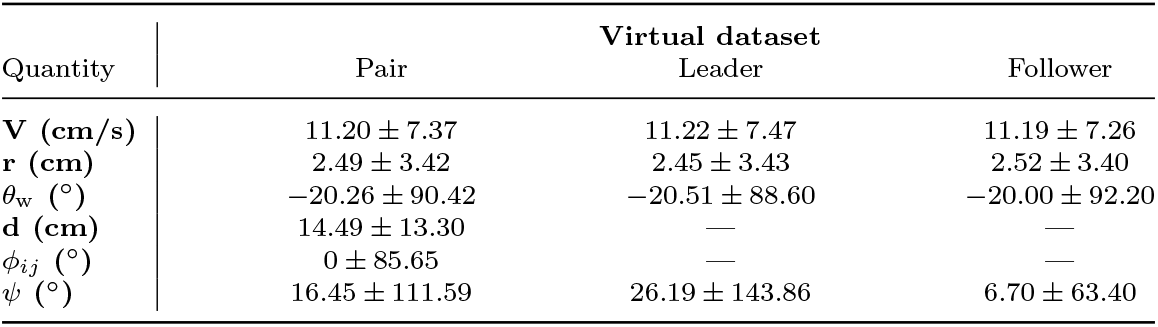
Mean and standard deviation values for *D. rerio*

**Supplementary Table 9:**
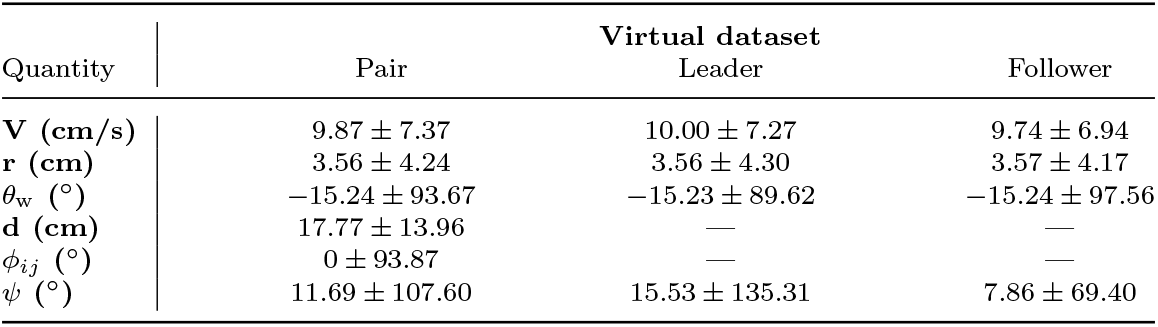
Mean and standard deviation values for DLI

## Footnotes

1 https://github.com/vita-epfl/trajnetplusplusbaselines

